# Total RNA sequencing reveals gene expression and microbial alterations shared by oral pre-malignant lesions and cancer

**DOI:** 10.1101/2023.03.24.534064

**Authors:** Mohammed Muzamil Khan, Jennifer Frustino, Alessandro Villa, Bach-Cuc Nguyen, Sook-Bin Woo, William Evan Johnson, Xaralabos Varelas, Maria Kukuruzinska, Stefano Monti

**Affiliations:** Section of Computational Biomedicine, Boston University Chobanian & Avedisian School of Medicine, Boston, MA, USA; Bioinformatics Graduate Program, Boston University, Boston, MA, USA; Department of Dentistry/Oral Oncology & Maxillofacial Prosthetics, Erie County Medical Center, Buffalo, NY, USA; Oral Medicine, Oral Oncology and Dentistry, Miami Cancer Institute, Baptist Health South Florida, Miami, FL, and Department of Orofacial Sciences, University of California San Francisco, San Francisco, CA, USA; Department of Translational Dental Medicine, Boston University School of Dental Medicine, Boston, MA, USA; Division of Oral Medicine and Dentistry, Brigham and Women’s Hospital and Harvard University, Boston, MA, USA; Division of Infectious Disease, Center for Data Science, Rutgers New Jersey Medical School, Newark, NJ, 07103, USA; Department of Biochemistry, Boston University Chobanian & Avedisian School of Medicine, Boston, MA, USA; Department of Biostatistics, Boston University School of Public Health, Boston, MA, USA

**Keywords:** Pre-malignant lesions, Oral cancer, oral microbiota, early detection

## Abstract

Head and neck cancers are a complex malignancy comprising multiple anatomical sites, with cancer of the oral cavity ranking among the deadliest and most disfiguring cancers globally. Oral cancer (OC) constitutes a subset of head and neck cancer cases, presenting primarily as tobacco-and alcohol-associated oral squamous cell carcinoma (OSCC), with a 5-year survival rate of ∼65%, partly due to the lack of early detection and effective treatments. OSCC arises from premalignant lesions (PMLs) in the oral cavity through a multi-step series of clinical and histopathological stages, including varying degrees of epithelial dysplasia. To gain insights into the molecular mechanisms associated with the progression of PMLs to OSCC, we profiled the whole transcriptome of 66 human PMLs comprising leukoplakia with dysplasia and hyperkeratosis non-reactive (HkNR) pathologies, alongside healthy controls and OSCC. Our data revealed that PMLs were enriched in gene signatures associated with cellular plasticity, such as partial EMT (p-EMT) phenotypes, and with immune response. Integrated analyses of the host transcriptome and microbiome further highlighted a significant association between differential microbial abundance and PML pathway activity, suggesting a contribution of the oral microbiome towards PML evolution to OSCC. Collectively, this study reveals molecular processes associated with PML progression that may help early diagnosis and disease interception at an early stage.

**AUTHOR SUMMARY:** Patients harboring oral premalignant lesions (PMLs) have an increased risk of developing oral squamous cell carcinoma (OSCC), but the underlying mechanisms driving transformation of PMLs to OSCC remain poorly understood. In this study, Khan et al., analyzed a newly generated dataset of gene expression and microbial profiles of oral tissues from patients diagnosed with PMLs from differing histopathological groups, including hyperkeratosis not reactive (*HkNR*) and dysplasia, comparing these profiles with OSCC and normal oral mucosa. Significant similarities between PMLs and OSCC were observed, with PMLs manifesting several cancer hallmarks, including oncogenic and immune pathways. The study also demonstrates associations between the abundance of multiple microbial species and PML groups, suggesting a potential contribution of the oral microbiome to the early stages of OSCC development. The study offers insights into the nature of the molecular, cellular and microbial heterogeneity of oral PMLs and suggests that molecular and clinical refinement of PMLs may provide opportunities for early disease detection and interception.

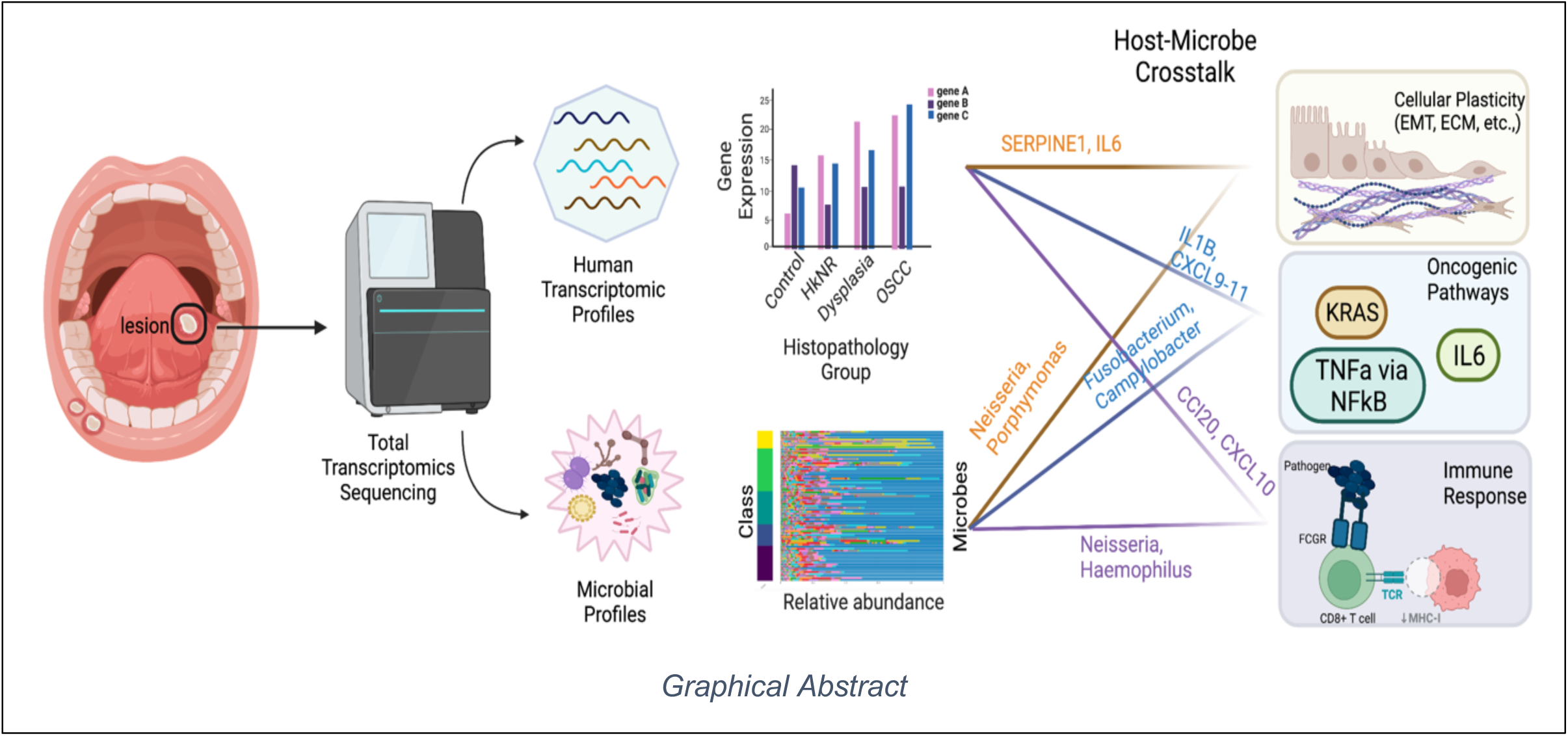

## INTRODUCTION

Head and neck cancer of the oral cavity ranks among the top ten most diagnosed cancers globally, with tobacco-and alcohol-associated oral squamous cell carcinoma (OSCC) pathology. OSCC arises from oral epithelial mucosa in a multi-step process in which normal cells are transformed into preneoplastic cells and then to cancer^1–3^. These premalignant lesions (PMLs) – also referred to as oral potentially malignant disorders (OPMD)^4^, but mostly presenting as leukoplakia and variants, and erythroplakia – transform to OSCCs through various histopathological stages, from hyperkeratosis/hyperplasia (or hyperkeratosis not reactive, *HkNR,* known previously as *keratosis of unknown significance(KUS)*), to various degrees of dysplasia^5^, carcinoma in situ (CIS), and finally to invasive OSCC^6^. The presence of dysplastic areas in the epithelium of the oral cavity has been associated with progression to cancer. Although oral dysplasia may progress to OSCC, the initial cause of their malignant transformation remains unclear. While many leukoplakias exhibit histopathology of dysplasia or invasive SCC at the time of biopsy, many non-dysplastic keratotic lesions (HkNR) also transform to invasive OSCCs over time, suggesting that such lesions may represent early dysplasia before the histologic phenotype of dysplasia is established^1, 3^. The genetic signatures of *leukoplakia* without dysplasia and other early oral lesions have not been well characterized. Thus, a key step to improving outcomes of OSCC is to identify the molecular factors driving disease initiation and progression, as these factors represent attractive candidates for disease interception through targeted therapies.

Here, we sought to uncover the mechanisms involved in OSCC development from PMLs by employing total RNA-sequencing and advanced computational analyses. A comprehensive knowledge of global alterations in gene expression programs and the detailed characterization of cancer-associated transcriptional and microbial signatures will provide new insights into the underlying molecular networks involved in OSCC initiation and progression, allowing the development of prognostic models of early OSCC pathobiology and the identification of markers for cancer risk assessment to improve prediction accuracy and disease outcome. Gene expression profiling represents a powerful tool to investigate progression in early oral dysplasia and HkNR^7, 8^. Importantly, adoption of a total RNA-sequencing technique allows us to simultaneously characterize the microbiota of the biopsies and to correlate microbial differences with host gene/pathway activity and clinical-pathologic features. While the role of the gut microbiome in inflammation and colorectal cancer has received much attention, little is known about the role of the oral microbiome in pre-malignant oral lesions^9^, and our study contributes to filling this knowledge gap.

## MATERIALS AND METHODS

### Subject population

A cohort of 66 individuals presenting with early oral lesions was chosen for this study (Table 1). Participants were enrolled prospectively from the Brigham and Women’s Hospital in Boston, MA, and Erie County Medical Center in Buffalo, NY, with protocols approved by the institutional review boards at the study sites. All study participants provided written informed consent. All patients underwent an oral biopsy and histopathological examination. The histopathology groups included normal mucosa (controls, n=18, 27%), hyperkeratosis not reactive (HkNR, n=17, 26%), moderate or severe dysplasia (dysplasia, n=22, 33%), and oral squamous cell carcinoma (OC, n=9, 14%). Smoking status for the missing patients were imputed using a random forest classification approach (see Data Analysis section below). In our study, the HkNR and dysplastic groups are collectively referred to as PML.

**Table 1:** Patient population stratified by histopathological group and clinical factors.

| Characteristic | Histopathology Group |  |  |  |
| --- | --- | --- | --- | --- |
|  | Control<br>N = 18 (27%)<sup>1</sup> | Hyperkeratotic; Not Reactive(HkNR)<br>N = 17 (26%)<sup>1</sup> | Dysplasia<br>N = 22 (33%)<sup>1</sup> | OSCC<br>N = 9 (14%)<sup>1</sup> |
| Age | 66 (51, 73) | 67 (30, 81) | 66 (44, 92) | 68 (49, 80) |
| Sex |  |  |  |  |
| Female | 11 (61%) | 11 (65%) | 11 (50%) | 5 (56%) |
| Male | 7 (39%) | 6 (35%) | 11 (50%) | 4 (44%) |
| Smoking_status |  |  |  |  |
| No | 11 (69%) | 7 (54%) | 10 (53%) | 7 (88%) |
| Yes | 5 (31%) | 6 (46%) | 9 (47%) | 1 (12%) |
| Unknown (Imputed) | 2 (No) | 4 (No) | 3 (No=1, Yes=2) | 1 (No) |
| Progression Status |  |  |  |  |
| Stable | 7 (70%) | 5 (71%) | 3 (38%) | 2 (100%) |
| Progressed to Dysplasia | 1 (10%) | 2 (29%) | 2 (25%) | 0 (0%) |
| Progressed to SCC | 2 (20%) | 0 (0%) | 3 (38%) | 0 (0%) |
| Unknown | 8 | 10 | 14 | 7 |

<sup>1</sup> Median (Range); n (%)

### Laboratory techniques

#### RNA sequencing

Total transcriptomic sequencing was performed on biopsy samples from 66 patients. Total RNA integrity was verified using RNA 6000 Pico Assay chips run on an Agilent 2100 Bioanalyzer (Agilent Technologies, Palo Alto, CA). To deplete ribosomal RNA, 100 ng of total RNA from each sample was incubated with RiboErase Mouse rRNA depletion oligos per the manufacturer’s instructions using Kapa RiboErase (HMR) Module (Kapa Biosystems, USA). Resulting rRNA duplexed with DNA oligos were digested with RNase H treatment. Libraries were then prepared using Kapa RNA HyperPrep Library

Preparation Kit according to the manufacturer’s protocol. Briefly, after rRNA depletion, the RNA was fragmented at 94C for 6 minutes. First strand cDNA was generated, and dUTP was incorporated during synthesis of second strand cDNA. Next, double-stranded cDNA underwent A-tailing, adaptor ligation, and incorporation of sample-specific multiplex indices before PCR amplification (12 cycles) according to the manufacturer’s protocol (Kapa Biosystems, USA). Size distribution and molarity of amplified cDNA libraries were assessed via the Bioanalyzer High Sensitivity DNA Assay (Agilent Technologies, USA). All cDNA libraries were sequenced on an Illumina NextSeq 500 instrument with 1.45 pM input and 1% PhiX control library spiked in (Illumina, USA) targeting 30-40 Million reads pairs per sample.

### Preprocessing and quality control

#### Human

The fastq files obtained from sequencing runs were aligned and mapped using *featureCounts()* from the Rsubread^10^ package. The human reference genome version used was GRCh38.99. Samples with a mapping quality of >40% were retained. Genes with counts greater than one count-per-million (cpm) in at least 9 samples (the lowest number of samples in a phenotypic group) were preserved and used for downstream analyses.

#### Microbiome

To obtain the metagenomic profiles, the same sequencing files that were used for human alignment and mapping were processed with Pathoscope 2.0^11^. The tool first removes all reads that align to the host transcriptome, with the remainder aligned and mapped to reference microbial taxonomies such as bacteria, fungi, and viruses. Bowtie alignment (the aligner used within Pathoscope) was run with parameters ‘`–local -k`’ (multiple local alignments option) set to 20, ‘`–min-score’` set to 90% (to extract only fragments that map to microbial genomes with high confidence) and read length, *L*, set to 70 (from fastQC). This yielded 948 operational taxonomical units (OTU) consisting of 323 genera across all samples. The whole analysis was carried out using the *animalcules* R package^12^

### Data analysis

#### Signature Projection

Several transcriptional signatures were used to interrogate our dataset (Figures 1C and 4). For each signature of interest, either paired sets of up-and down-regulated genes, or a one-directional geneset were defined. For each geneset, an aggregate “activity” score was computed by *Gene Set Variation Analysis* (GSVA)^13^. For bidirectional signatures, a combined score was computed based on the difference of the up-and down-regulated GSVA scores. The following signatures were derived from publicly available datasets: 1) a signature of the comparison of Adjacent Epithelial (AE, n=44) with Head and Neck Squamous Cell Carcinoma (HNSCC, n=500) from the TCGA^14^, consisting of 1317 up-and 1330 down-regulated markers in HNSC; 2) a signature comparing malignant-transforming (MT, n=10) with not-transforming (NT, n=10) *oral potentially malignant disorders* (OPMD)^15^, consisting of 9 up-and 28 down-regulated markers. OPMD were there defined as oral lesions diagnosed as leukoplakia or erythroleukoplakia, and carrying a potential risk of malignant transformation^15^; 3) a one-directional signature of “partial epithelial-to-mesenchymal transition” (pEMT), a molecular phenotype characteristic of cells spatially localized to the leading edge of primary tumors and shown to be an independent predictor of nodal metastasis^16^.

**Figure 1:**
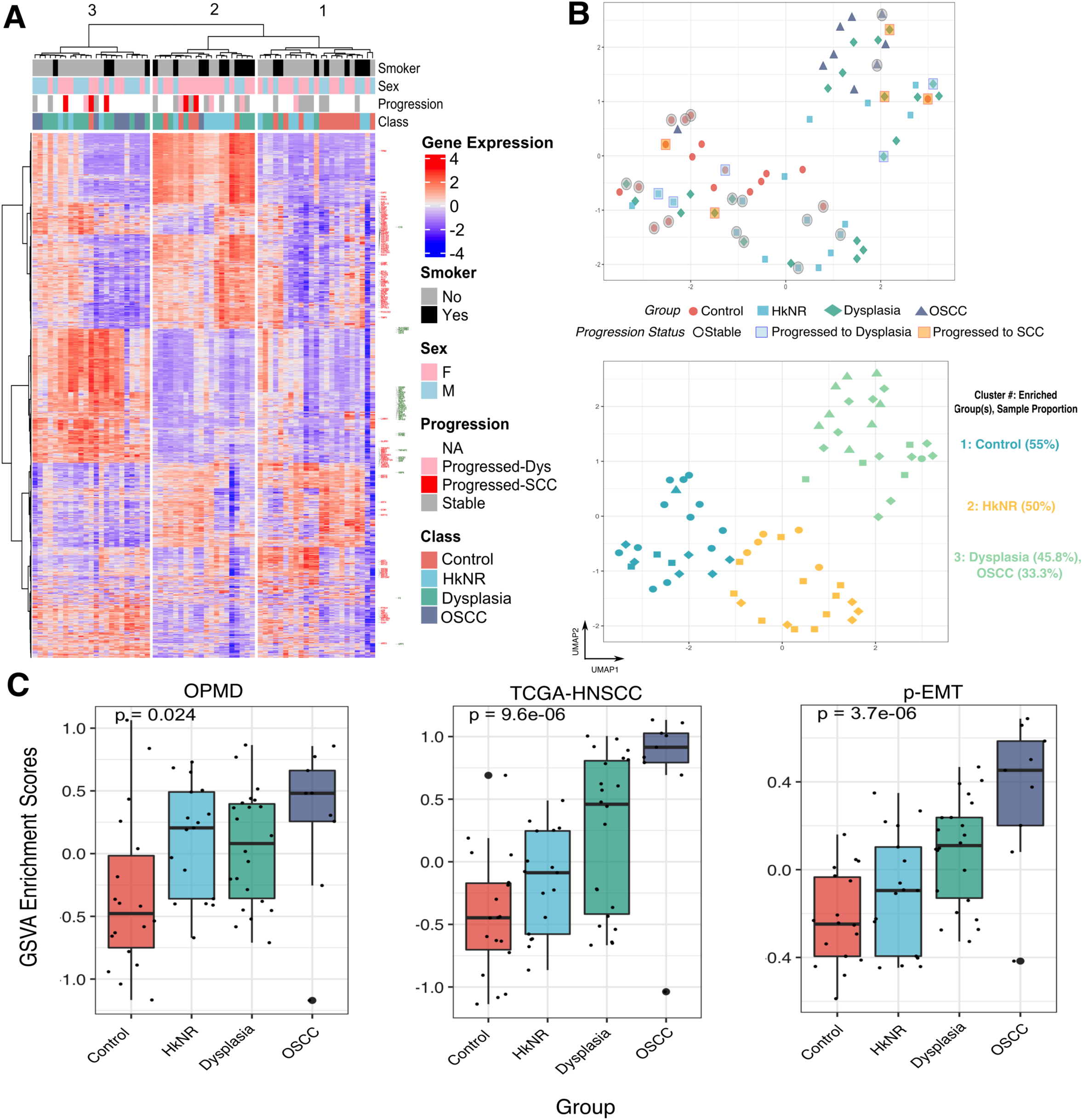
PMLs show transcriptional heterogeneity. Figure 1A: Heatmap of top 2000 highly varying genes, with rows (genes) and columns (patients) sorted by hierarchical clustering. The columns are split into three groups as determined by a 3-way cut of the dendrogram. Row annotation highlights epithelium-related genes in red and immune-related genes in green. Figure 1B (top): UMAP analysis of gene expression profiles from all samples colored by histopathological group. The progression status of 27 patients is denoted with black circles (stable), blue squares (progressed to dysplasia), and red squares (progressed to SCC). Figure 1B (bottom): Clustering analysis using K-means, with k=3. Within each cluster, the group positively enriched by chi-square test is highlighted. Figure 1C: Boxplots of GSVA-based enrichment scores. Displayed from left to right are: score differences of up-and down-regulated OPMD signatures (n=37 genes; up=9, down=27); score differences of up-and down-regulated TCGA HNSCC signatures (n=2647; up=1317, down=1330); and scores from a unidirectional p-EMT signature (n=100).

#### Dimensionality reduction and visualization

UMAP dimensionality reduction analysis was performed based on the 2^nd^ through 10^th^ principal components (PC) derived from the top 500 highly variable genes in the dataset. Probabilistic clustering as implemented in the *mclust*^17^ package was performed to estimate the number of clusters, which yielded k=3 clusters, followed by application of k-means clustering, which yielded the cluster assignments depicted in Figure 1B. Cluster enrichment for any of the lesion groups was assessed based on a three-group *chi-square* test, with the group labeling a cluster identified as the one with a positive and higher chi-square score than the other groups.

#### Smoking Status imputation

Smoking status, when missing, was estimated based on imputation. To this end, multiple classifiers – random forest (RF), support vector machine (SVM), and K nearest neighbors (KNN) – were evaluated by 10-fold cross validation on the dataset restricted to the top 1000 genes with highest median absolute deviation (MAD) and the samples with available smoking status, with the RF classifier yielding the best area under the curve (AUC = 0.79). A RF classifier trained on all complete data (n=56) was then applied to the prediction of the missing smoking status (n=10).

#### Differential analysis

Differential gene expression analysis was performed based on DESeq2^18^. Age, sex, and smoking status were included as covariates in the analysis. Differential signatures were defined as the sets of transcripts with FDR-corrected q-value≤0.05 and log_2_ fold-change≥1.5 for up-regulated signatures and log_2_ fold-change ≤ -1.5 for down-regulated signatures.

#### Pathway enrichment analysis

Differential signatures were annotated by over-representation analysis (ORA) based on hyper-geometric test using the hypeR package^19^ and the HALLMARK and REACTOME geneset compendia available through MSigDB^20, 21^. hypeR was run with the complete set of genes in the dataset as background (n=21,500), and the significant pathways were defined as those with an FDR-corrected q-value≤0.1. The hierarchy of pathways as depicted in Figure 2B was derived using the hierarchicalSets^22^ method.

**Figure 2:**
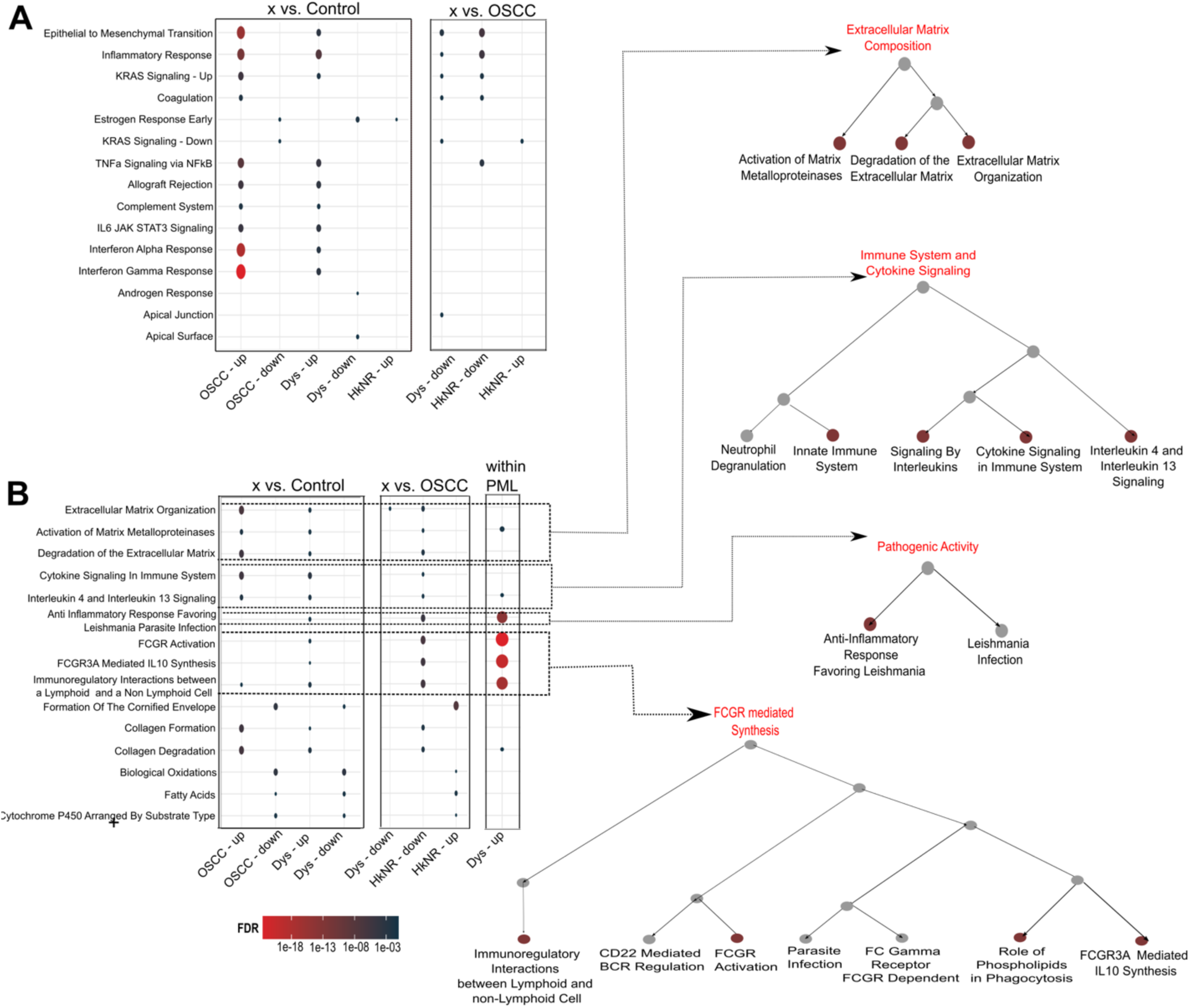
Pathway enrichment of the PML and OSCC signatures. Figure 2A: Over-representation-based enrichment results based on the Hallmarks compendium of differentially expressed genes from pairwise analysis with control, OSCC, and within PMLs. The size of the dot is proportional to the number of overlapping genes and the color coding represents the FDR-corrected q-val. Figure 2B: Over-representation-based enrichment results based on the Reactome compendium. The hierarchical organization of related pathways is shown on the right.

#### Immune cell type deconvolution

Using TPM-normalized (transcripts per million) counts, *CIBERSORT*^23^ was applied with reference signatures from immune cell types derived from human hematopoietic cells consisting of 22 main immune cell-types.

#### Microbiome analysis

The microbiome analysis was carried out using *animalcules*^12^, which provides various functions to create a multi-assay experiment object that contains the relative abundance of microbes detected in the study and several visualization methods. The relative abundance and diversity plots in Supplementary Figures S5 A-C were generated using this package. DESeq2-based^18^ differential analysis of microbes abundance between groups was also performed using an *animalcules* wrapper. An FDR-corrected q-value≤0.05 was used to select differentially abundant microbes.

#### Microbe-set enrichment analysis and pathway activation

The microbe-set enrichment analysis (MSEA)^24^ python package was used to establish significant associations between microbes and genes. MSEA defines gene-labeled microbe-sets by tallying gene-microbe co-occurrences in PUBMED publications, and by then grouping microbes that co-occur with the same gene. For example, the microbe-set for gene UHRF2 consists of the 4 microbes (*Porphymonas, Haemophilus, Fusobacterium, and Campylobacter*) found to “significantly” co-occur with that gene across publications based on a Jaccard index criteria. A library of ∼1,300 microbe-sets thus defined, corresponding to as many genes, was used for our analysis^24^.

Next, microbial genera found to be differentially abundant in OSCC and PML compared to the control group were tested for enrichment against this microbe-set library, and the results visualized as a bipartite graph consisting of microbe-gene interactions with a significant FDR-corrected q-value≤0.05 and a positive MSEA combined score.

Finally, the union of mset-labeling genes was tested for enrichment of the significant Hallmark genesets found in the host analyses (OSCC *vs.* control and PML *vs.* control, Figure 2A). Genes in the overlap between Hallmarks and the host differential signatures were tested against the mset-labeling genes, and those significant at a q≤0.25 were reported.

## RESULTS

### Patient Population

A cohort of 66 individuals selected for this study included patients with oral lesions representing controls (normal mucosa, 27%), hyperkeratosis not reactive (HkNR, 26%), moderate or severe dysplasia (dysplasia, 33%), and oral squamous cell carcinoma (OSCC, 14%) (Table 1). The mean age of patients was 55. Clinical characteristics such as smoking status – current or former, and follow-up information on the progression of the lesions were included, when available. Progression status for 27 of the 66 patients was available, with 10 patients progressing to the next stage of pre-cancer or OSCC pathology. In particular, 2 patients with HkNR histopathology transformed to dysplasia, and 3 patients with dysplastic phenotype progressed to OSCC, accounting for ∼8% of the PML population. Additionally, 2 patients in the control group progressed to OSCC.

### The transcriptional profiles of PMLs display heterogeneity

We selected the top 2000 highly variable genes based on median absolute deviation (MAD) and visualized their expression profiles on a heatmap in Figure 1A, with the rows (genes) and columns (samples) clustered across histopathological groups in an unsupervised manner. The heatmap was partitioned into the three top branches identified by hierarchical clustering, which was independent of the probabilistic clustering results shown in Figure 1B (bottom panel). Importantly, the groupings displayed in Figure 1A and 1B captured similar salient features of the histopathological groups, attesting to the robustness of the transcriptionally driven patient segregation. In particular, Cluster 1 predominantly contained control samples, cluster 2 consisted primarily of PMLs and some of the control samples, and cluster 3 included all but one OSCC samples along with some PMLs. Of note, cluster 3 also contained the control sample that progressed to OSCC. Several genes within pathways that were the focus of subsequent sections are highlighted. For example, genes associated with epithelial and extracellular matrix-related functions, such as COL7A1, TGFB1, LAMA3, and KRT10, are labeled in red, and immune-related genes, such as HLA-A, HLA-B, and STAT1, are labeled in green. From this heatmap a pattern emerged where all the OSCC samples and some of the dysplastic and HkNR samples showed high expression in most of the genes related to these processes, with the remaining samples showing patterns that matched the control group. Importantly, some of the samples from the non-cancer groups that progressed to OSCC showed transcriptional patterns similar to those of OSCC.

While the heatmap shows transcriptional heterogeneity, the unsupervised organization of the PML groups determined using a UMAP dimensionality reduction approach showed the clear separation of the control and OSCC groups, with the PML groups distributed along a gradient between the two extremes (Figure 1B, top panel). Within these groups, most of the dysplastic samples were located proximal to the OSCC group region on the upper right side of the plot, while the HkNR samples were more scattered in-between suggesting more heterogeneous phenotypes. Model-based probabilistic clustering^17^ was applied to estimate in an unbiased, data-driven fashion the number of clusters and their composition, which yielded a 3-cluster partition. The resulting cluster assignments are shown in Figure 1B (bottom panel), with cluster 1 comprising a majority of control samples, cluster 2 comprising some control samples and a significant proportion of HkNR, and cluster 3 including all the OSCC samples and a significant proportion of the dysplastic samples.

Figure 1B (bottom panel) also shows the progression status of patients (n=27) obtained through follow-up visits. Progression status is denoted by red and blue shaded boxes, while black denotes a stable condition. Patients with dysplasia that transformed to OSCC segregated towards the cancer cluster (cluster 3). Of note, a control sample that progressed to OSCC was clustered with the cancer group (cluster 3) by the UMAP analysis, suggesting clinical relevance of the associated molecular profile. This sample had a unique transcriptional profile that shared similarities with the SCC group as shown in Figure 1A.

Taken together, these unsupervised analyses highlight the considerable transcriptional heterogeneity of PMLs. The limited availability of lesions with long-term follow-up precluded us from drawing stronger conclusions, but we note that PMLs likely to progress share greater transcriptional similarity with OSCCs than non-progressing PMLs.

### PMLs are enriched for malignant-transforming and partial-EMT signatures

To evaluate the transcriptional patterns associated with progressive malignant transformation in our datasets, we leveraged multiple signatures from published studies capturing salient features of such a transformation. These included: a tumor *vs.* normal signature derived from the cancer genome atlas (TCGA) of HNSCC RNAseq dataset^14^; a signature of malignant-transforming (MT) *vs.* not-transforming (NT) OPMD^15^ ; and a signature of “partial-EMT” (pEMT) – a molecular phenotype shown to be an independent predictor of nodal metastasis^16^. For each of the signatures, GSVA-based enrichment scores corresponding to the up-and down-regulated genes were estimated for each of the samples, and the difference scores (up – down) were then computed. The scores were then stratified by histopathology group (Control, HkNR, Dysplasia, OSCC), as shown in Figure 1C.

When stratifying the samples by the OPMD signature, HkNR showed higher and more variable enrichment scores amongst the PML. Stratification by the TCGA HNSCC “tumor *vs.* normal” signature showed a clear upward trend tracking with progression, as did stratification by the p-EMT signature, a defining characteristic of more aggressive HNSCs^16^.

In summary, these results confirm that our data capture salient features of transformation identified in previous studies and help further refine the transcriptional programs shared by different oral lesions. In particular, known signatures of premalignant and malignant transformation and tumor aggressiveness tracked with the canonical lesion progression represented by our histopathology groups, with interesting exceptions. Notably, the HkNR group displayed a higher level of the OPMD signature (but also higher variability) than the putatively more advanced dysplasia.

### PML signatures are associated with oncogenic and immune-altering pathways

The observed highly varying transcriptional profiles of PMLs (Figure 1A-B) may contribute to current challenges in their evaluation and clinical treatment even with histopathology information. To gain further insight into the transcriptional programs defining the different groups, we performed pairwise differential gene expression and pathway analyses using *DESeq2* and *hypeR*, respectively. Differential analysis was performed by comparing each remaining group to controls, to OSCC, and by pairwise comparing the PML sub-groups, while controlling for sex and smoking status. The number of significant markers along with differential analysis results for each pairwise comparison is reported in the Supplementary section (Figure S1 and Supplementary Tables 1-3). As shown in Figure 2, the signature genes with a *logFC* cutoff of ±1.5 and q-value ≤ 0.05 were found to be significantly enriched for several immune response pathways defined in the Hallmark and Reactome compendia. Hallmark pathways that were significantly enriched in the OSCC and dysplastic groups compared to controls included key cancer progression and transformation pathways, such as epithelial-to-mesenchymal transition (EMT) and KRAS signaling, as well as inflammatory response pathways such as TNF-α signaling via NF-κB, interferon gamma (IFN-γ) and alpha (IFN-α) response, IL2/STAT5, and IL6/JAK-STAT3 signaling. Genes driving the EMT enrichment in both groups included extracellular matrix-related genes such as ADAM12, TGFB1, LAMA3, MMP1 and MMP3, and in the OSCC group multiple collagen genes (COL11A1, COL12A1, COL4A1) and serine proteinase inhibitors SERPINE1 and SERPINH1, among others, while genes driving the IFN-γ and IFN-α enrichment in both groups included multiple chemokine ligands (CXCL9, CXCL10, CXCL11), and in the OSCC group multiple interferon induced proteins (IFIT1-3, IFI35, IFI44, etc.) among others (see Supplementary Table 2). The comparisons to the cancer group revealed similar enrichment patterns of immune response pathways in the cancer group compared to the dysplastic and the HkNR groups.

The differential signatures were then tested for enrichment with pathways in the Reactome compendium to evaluate potential metabolic and pathogenic associations. Multiple pathways from the same biological processes were found to be enriched and were thus organized and visualized with their hierarchical structure in Figure 2B. This analysis showed a significant enrichment, in PMLs and OSCCs when compared to control, of tumor microenvironment-related activities, such as extracellular matrix (ECM) organization, and activation of matrix metalloproteinases. Confirming the Hallmarks-based analysis, the Reactome-based analysis also identified and further resolved several enrichments of immune and pathogenic pathways. Signaling by interleukins, the FC gamma receptor (FCGR) mediated regulation, and FCGR phagocytosis were enriched in the OSCC and dysplasia groups when compared to controls and HkNR groups. Figure 2B shows their hierarchical organization, with multiple FCGR-related pathways, as well as multiple immune-related pathways.

The strong enrichment of p-EMT and EMT signatures in the PML groups raised the question of whether this enrichment stems from increased plasticity within the epithelial compartment or from alterations in adjacent stroma. To investigate this, we tested our series for enrichment with respect to a cancer-associated fibroblast (CAF) signature previously described^25^, inclusive of genes COL1A1, COL1A2, COL3A1, and PDGFRB, among others. Interestingly, the signature was significantly enriched in the non-control groups, with the strongest enrichment observed in the PML groups (Supplementary Figure S2B). Furthermore, CAF enrichment was significantly positively associated with p-EMT enrichment (Supplementary Figure S2C), with the association stronger in PMLs. These observations suggest that CAF infiltration and increased plasticity co-occur in PMLs, and single cell-based studies will be needed to further elucidate their interplay.

### PMLs and OSCC have significantly higher levels of immune activity than controls

Our differential gene and pathway analyses showed that the PMLs shared significant similarities with the OSCC group. Given the observed up-regulation of multiple immune pathways, we next focused on the immune landscape by investigating changes in immune cell proportions across healthy and disease groups. To this end, we performed immune cell type deconvolution using a gene signature compendium comprising 22 immune cell-types derived from human hematopoietic cells. The cell proportions estimated with CIBERSORT, visualized as boxplots in Figure 3, were stratified by histopathology group at different levels of resolution. Immune cell types were first divided into the innate and adaptive immune components in the top panel of Figure 3. These were further subdivided into their main subtypes, with the innate component subdivided into macrophages (M0-like, M1-like and M2-like), monocytes, mast cells, NK cells and neutrophils (Figure 3, bottom left panel), and the adaptive immune component subdivided into B-and T-cells (Figure 3, bottom right panel). Of note, the abundance of all innate immune cell subtypes significantly increased in non-control groups, including PMLs and cancer groups. Also, B-and T-cells showed an increase in non-control groups but did not reach significance. A detailed breakdown of the cellular activities in these immune cells is shown in Supplementary Figure S4 A-B. For example, CD8+ T-cells, Tregs and CD4 memory resting/activated cells showed an increased abundance in PMLs and cancer. Sample level breakdown is visualized as a heatmap in Supplementary Figure S4C.

**Figure 3:**
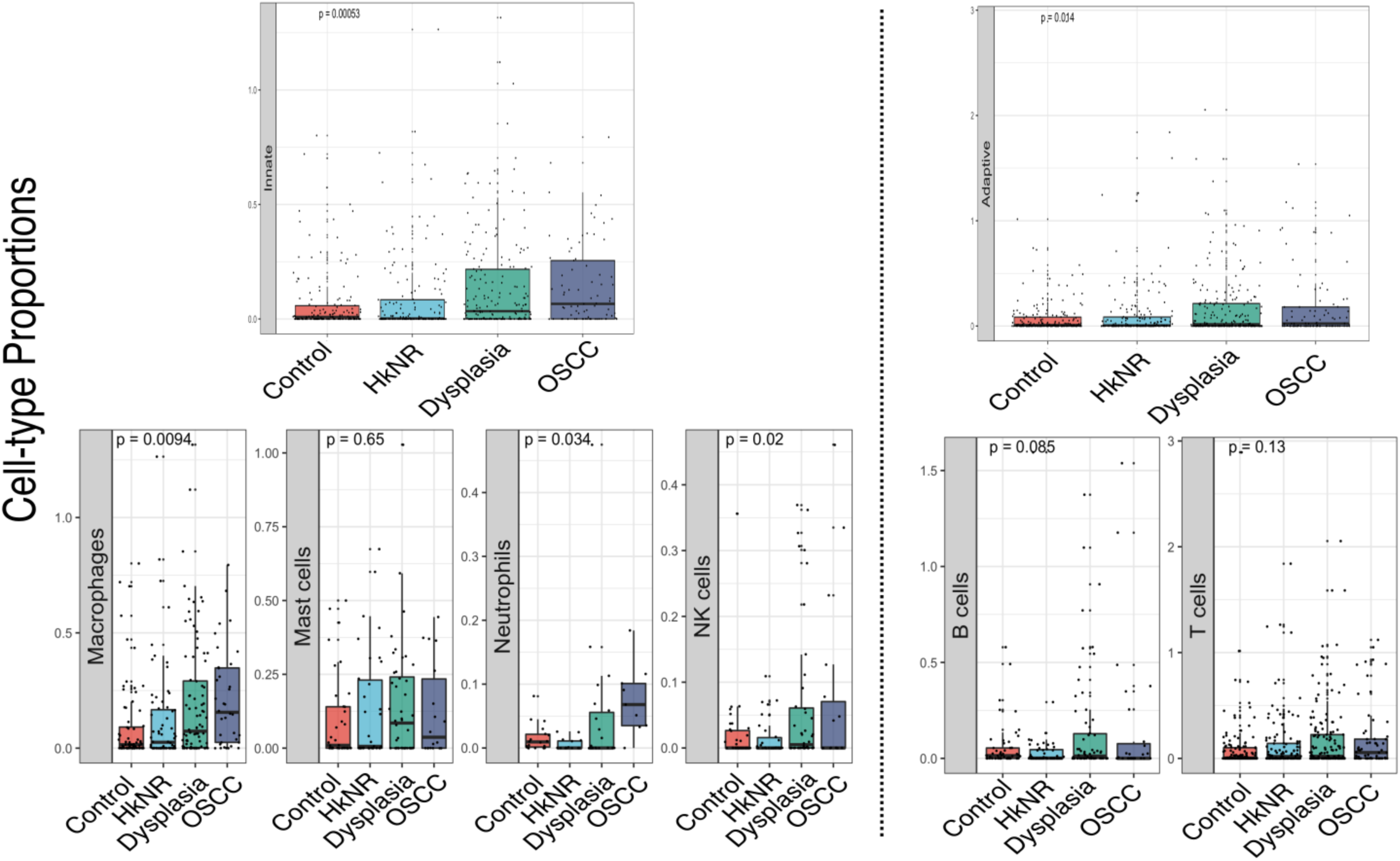
Relative abundance of immune cell-types stratified by histopathology. Immune cell-type deconvolution scores obtained from CIBERSORT are stratified by histopathological groups. (Top) Cell type proportions of the innate (left) and adaptive (right) compartments. (Bottom) Specific cell sub-type proportions within the two compartments.

### Oral PMLs show transcriptional similarities with bronchial PMLs

Given the contiguity of the oral cavity in the upper airway with the lungs in the lower airway, we next assessed the transcriptional similarities between our oral PMLs and bronchial PMLs. Beane et al.^26^ identified four distinct PML bronchial subgroups, including a proliferative and an inflammatory response sub-groups, and nine gene co-expression modules recapitulating their transcriptional programs. Significantly, the gene modules corresponding to cell-cycle (proliferative sub-group), inflammatory response and IFN signaling (inflammatory response sub-group) showed enrichment in our non-control groups (Supplementary Figure S3), suggesting that oral lesions from the upper airway region and lesions from the main airway compartment share common processes likely to be implicated in their progression to malignancy.

### Differentially abundant microbes in PMLs and OSCC are associated with pathways that promote malignancy and immune activity

The relative abundance of the microbial taxa stratified by the phylum, genus and species levels are plotted in Figure 4A. Overall, the microbial abundance across groups varied, with the genus of Fusobacterium dominating the OSCC group, and Streptococcus prominent in the PML group. The alpha and beta diversity stratified by the histopathological group were higher in the OSCC and PML groups compared to the healthy individuals (Supplementary Figure S5 A-B). To identify microbes (at the genus level) that were differentially abundant in PML and OSCC compared to the control group, we performed differential abundance analysis. With an FDR-corrected q-value ≤ 0.05, 27 microbes were more abundant in the OSCC group than the control group, 8 in HkNR and 28 in dysplasia (Supplementary Table 4). Some of the microbial genera that were significantly more abundant in both PML and OSCC groups than in the control group were *Neisseria, Prevotella,* and *Fusobacterium* (Supplementary Figure S5C). These have all been shown to be associated with cancer in previous studies^27–31^.

**Figure 4:**
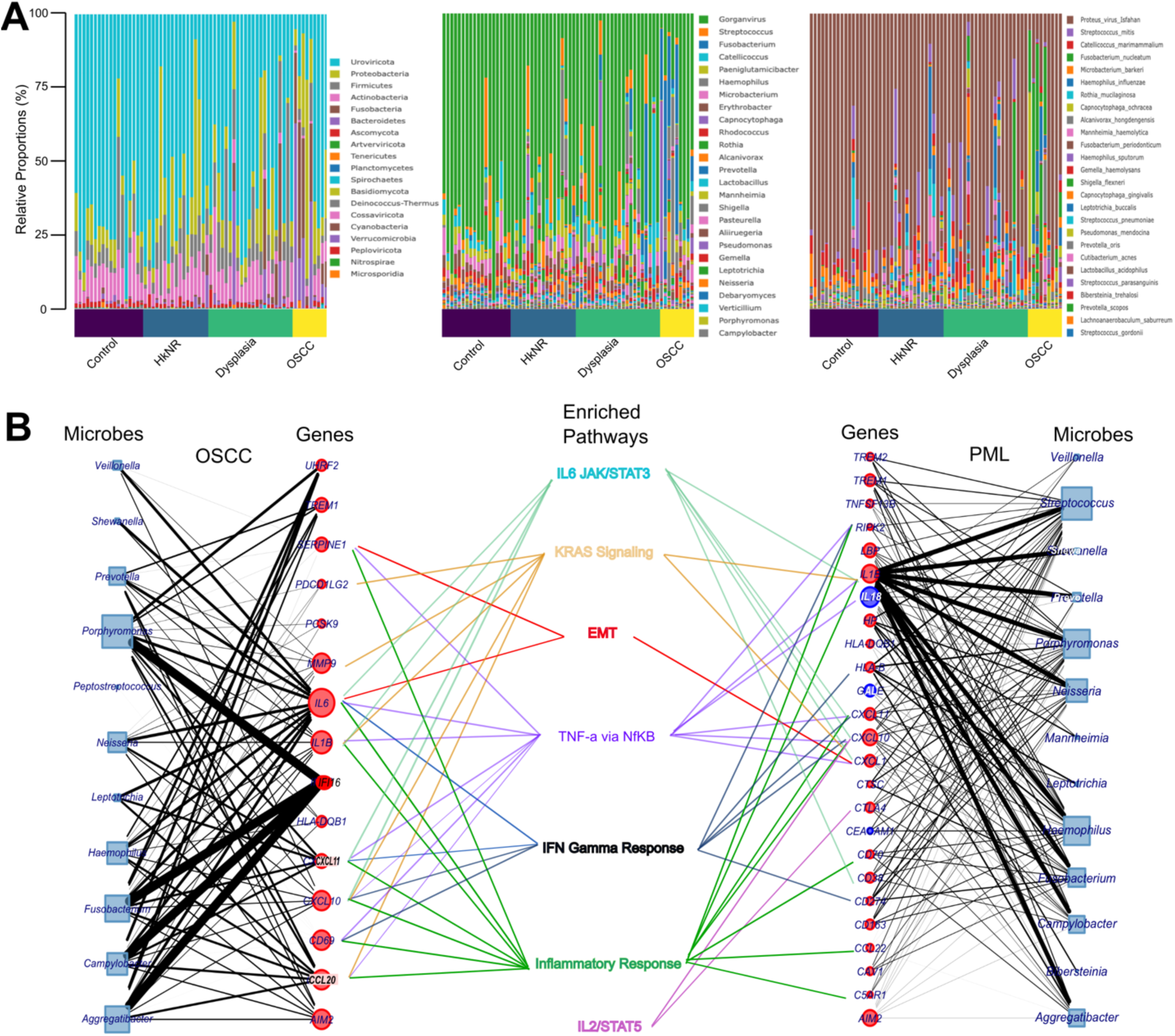
Differential microbial abundance and their association with PML and OSCC. Figure 4A: Log-CPM normalized relative proportions of microbial profiles of Phylum, Genus and Species level stratified by histopathological grouping. Figure 4B: Associations of differentially abundant microbes with human genes and host-response pathways are outlined. The node size denotes the total number of associations for a microbe or a gene, and the edges’ thickness is proportional to the enrichment score between a microbe and a gene. The node color indicates whether the gene was up-(red) or down-regulated (blue) in the corresponding host transcriptome differential analysis. The gene-pathway edges denote enrichment by over-representation analysis.

To explore potential associations between differentially abundant microbial genera and the host transcriptome, we performed microbe-set enrichment analysis (MSEA)^24^, whereby the differentially abundant microbes we identified were tested for enrichment against a library of ∼1,300 gene-labeled microbe-sets, and the significant microbe-gene associations were visualized in Figure 4 and tabulated in Supplementary Table 5. The union set of genes labeling significantly enriched microbe-sets was in turn tested for enrichment against the Hallmark compendium using hypeR (Supplementary Table 6) The significant genesets overlapping with those found in the host analysis in Figure 2A are also displayed in Figure 4 (center).

The genesets corresponding to EMT, KRAS signaling, IL6/STAT/JAK3, IL2/STAT5, IFN Gamma Response, Inflammatory Response, and TNFa via NFkB, were found to be significantly enriched in both analyses, suggesting a potential mediation by the microbiome of the association between OSCC’s and PML’s phenotypes and selected cancer-and immune-related pathways. For example, when comparing OSCC to control, *Veillonella*, *Aggregatibacter*, *Prevotella*, *Porphyromonas*, and *Neisseria* drove the enrichment with respect to both the SERPINE1-and IL6-indexed microbe-sets, which in turn are included in the Hallmarks’ EMT geneset (Supplementary Table 6). Similarly, *Fusobacterium*, *Campylobacter*, *Aggregatibacter* and *Porphymonas* drove the enrichment with respect to the IFI16-indexed microbe-set. IFI16 is an interferon gamma inducible gene that activates the pro-inflammatory pathway IFN-ξ. Differentially abundant microbes were also enriched for the microbe-sets labeled by CXCL10, CXCL11, and CCL20, among others. Similar patterns were observed for the PML group.

## DISCUSSION

OSCC are thought to arise from PMLs in the oral cavity, which have the potential to transform to cancer through a multi-step series of clinical and histopathological stages. Histologically, PMLs may represent varying degrees of oral epithelial dysplasia, although very little is known about pre-OSCC biology^1^. Characterization of the transcriptional programs defining these premalignant stages may help identify important mechanisms of early transformation and support the design of novel interception strategies. To this end, we generated total RNA sequencing profiles from biopsy samples of 66 patients comprising healthy controls, PML lesions that included HkNR and dysplasia, as well as OSCC, with the goal of defining molecular profiles associated with dysplasia pathobiology. We performed statistical and bioinformatics analysis of the transcriptional profiles to define and annotate signatures corresponding to the distinct patient groups, and to investigate the biological pathways associated with progressive transformation. Importantly, availability of the total transcriptome allowed us to query and overlay the oral microbial gene signatures onto the host transcriptome.

Our results indicate that PMLs show considerable heterogeneity, as highlighted in Figure 1. Of particular interest was the large heterogeneity manifested by the HkNR group based on hierarchical clustering as well as on dimensionality reduction and model-based clustering analyses (Figure 1A-C), with subjects from that group included in all three clusters (although predominantly in the middle cluster). HkNR has been recognized as an early dysplastic lesion with significant malignant transformation potential^35^, and its transcriptional heterogeneity described here suggests the need for a molecular refinement of this broad phenotype. The availability of follow-up information for a subset of the patients, and their segregation tracking with progression placement in the control-to-cancer gradient (Figure 1B) also suggests the feasibility of developing prognostic biomarkers based on expression profiles. Still, the latter will require a larger PML patient cohort than the one available in this study.

Differential signature and pathway analyses across the different groups showed a significant early activation of cancer-associated pathways (EMT and KRAS signaling) as well as of immune and inflammatory pathways, confirming the important role played by the immune system in malignant transformation^32, 33^. Further mechanistic studies will be necessary to elucidate whether the activation of these pathways is a response to, or driver of, such transformation. It should be emphasized that, within the PML group, these enrichments were mainly found in the dysplastic samples, reflecting the higher heterogeneity of the HkNR sub-group. Also relevant was a significant enrichment of Fc gamma receptor (FCGR)-mediated regulation and FCGR phagocytosis in OSCC and dysplasia compared to control. Fc gamma receptors are a family of heterogeneous molecules that play both activating and inhibitory tumor activity, and can bind to immunoglobulin (Ig) classes and subclasses, including those present in infected cells and pathogens^34^. Once bound, the FCGR phagocytosis process functions to eliminate the unwanted pathogenic activity^35, 36^. This may suggest that to prevent PMLs from progressing to malignancy, these inflammatory pathways are recruited to remove any of the pathogens or abnormal cells.

Another relevant finding was the enrichment in cancer and PMLs in pathways associated with extracellular matrix (ECM) organization, and activation of matrix metalloproteinases. The former has known roles in cell proliferation and migration, and the latter has a role in wound healing but adverse effects in cancer when activated by certain growth factors^37^.

The oral microbiota is the second most diverse microflora in the human body after the gut. The relative abundance of several microbial genera in Supplementary Figure S5 is an indication of how varied the microbiota of the PML groups is when compared to the normal oral mucosa (control group), with a higher alpha diversity measure (Supplementary Figure S5A). Varying levels of certain microbes can be correlated with disease progression, and our results confirm this association. For instance, *Fusobacterium* – a gram-negative bacterium significantly more abundant in PML and OSCC than in control in our series (Figure S5C and Supplementary Table 2) – has been shown to be abundant in patients diagnosed with oral cancer^27, 31^. *Fusobacterium nucleatum*, a sub-species of the *Fusobacterium* genus, has adhesion properties such that it may latch onto other bacteria and infected cells and promote an EMT-like phenotype and metastasis in some cases^28^. This role was supported by our MSEA analysis which showed a significant association with the EMT phenotype (Figure 4B). Similarly, the genus of *Neisseria*, also more abundant in PML and OSCC than in control in our series (Figure S5C and Supplementary Table 2), has been hypothesized to play a role in alcohol-related oral carcinogenesis. Taken together, our results support the conclusion that an increase in the activity of these microbes is associated with PML progression^30^, and functional studies will be needed to decode the details of the observed associations.

In summary, our results indicate that higher levels of gene expression in PMLs and OSCCs are associated with selected oncogenic pathways, including EMT and KRAS signaling, as well as with pro-and anti-inflammatory pathways, which in turn are associated with the differential abundance of multiple microbial genera^38^.

## ACKNOWLEDGEMENTS

We acknowledge the TCGA Research Network (https://www.cancer.gov/tcga) for granting access to cancer patient bulk expression data. We would like to thank David Sherr and Manish Bais for their feedback on the interpretation of the study results.

## FUNDING

This study was supported by NIH/NIDCR grants 5 R01 DE030350 (MAK, SM, XV), R01 DE031831 (SM), NIH/NCATS grant BU-CTSI 1UL1TR001430 (SM), ACS Research Scholar Award RSG-17-138-01-CSM (XV), Research for Health in Erie County, Inc. (JF), and R01 GM127430 and R21 AI154387 (WEJ). Its contents are solely the responsibility of the authors and do not necessarily represent the official views of the NIH. All authors have read the journal’s authorship agreement and policy on disclosure of potential conflicts of interest.

## COMPETING INTERESTS

The authors declare no competing interests.

## DATA & CODE AVAILABILITY

The RNA-sequencing data of the 66 samples here described are available at Gene Expression Omnibus (GEO), accession ***GSE227919*** (provisional release date: 12/31/2023, which will be updated upon publication). The scripts used to perform the analysis can be accessed at https://github.com/montilab/pml_analysis.

## Supplementary Figures

**S1.**
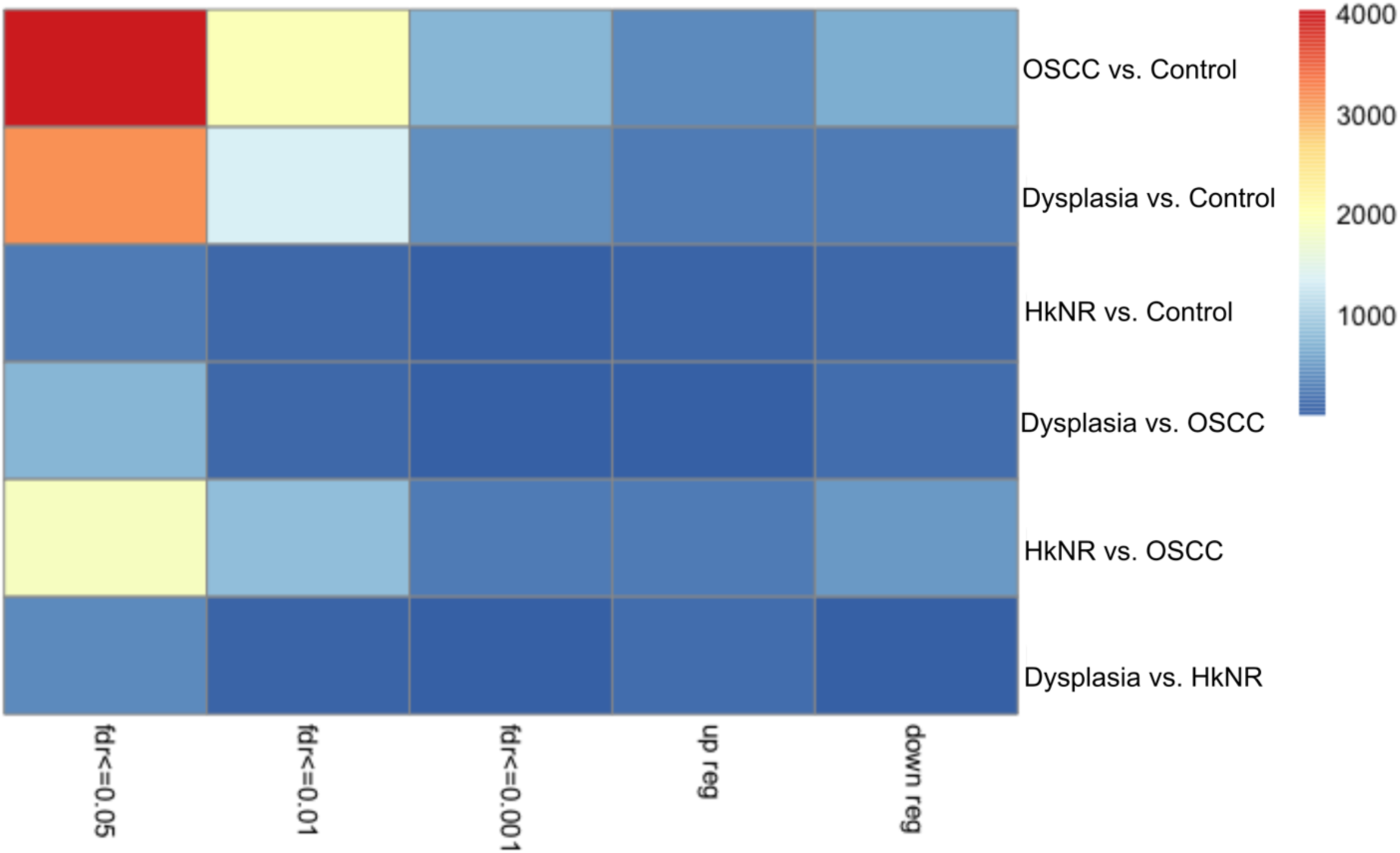
Total number of markers per pairwise analysis. Total number of differentially expressed genes in the pairwise analysis with logFC ± 1.5 and q-val :-s {0.01, 0.05, 0.1}.

**S2.**
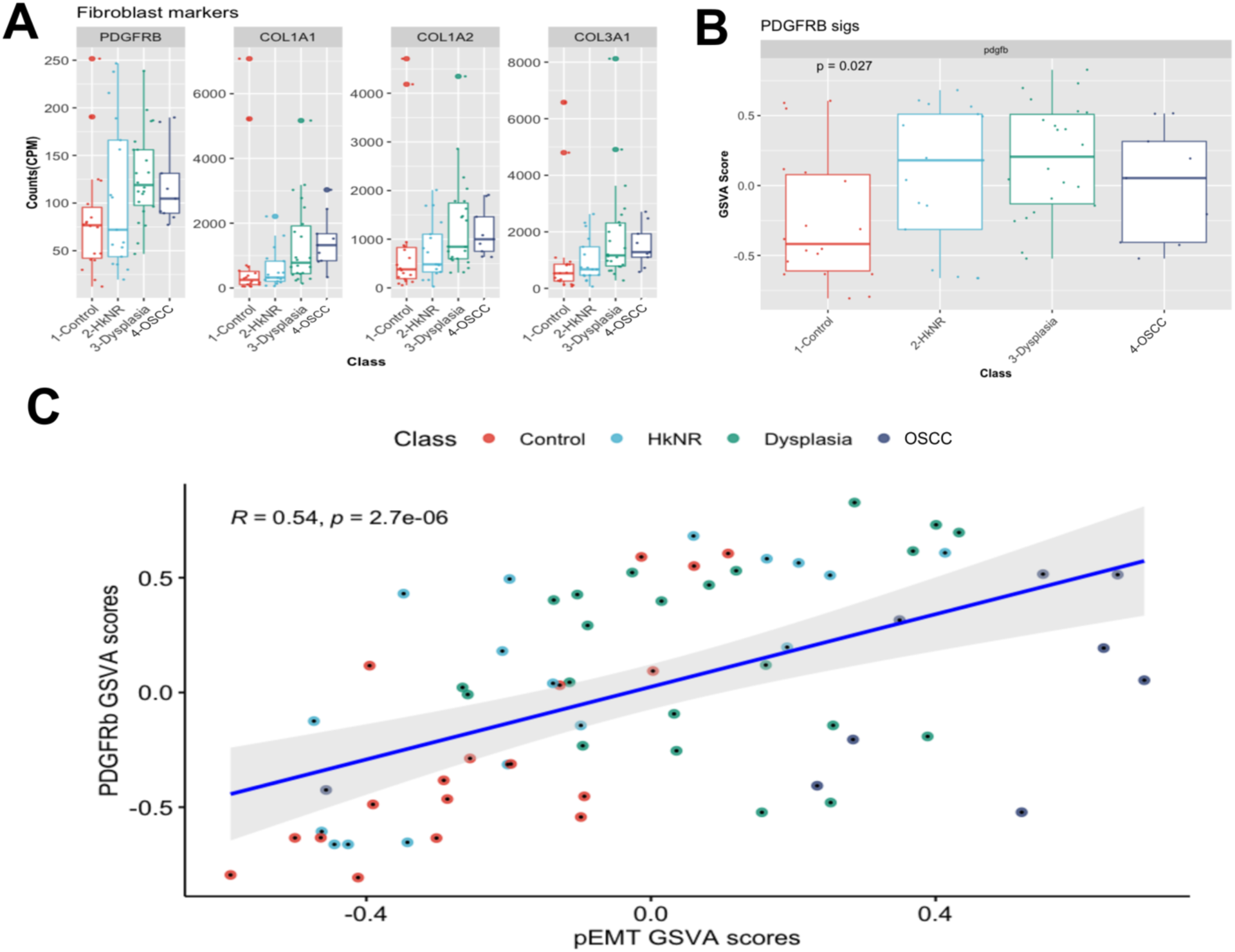
Enrichment of cancer-associated fibroblasts. A. Gene expression of PDGFB1, COL1A1, COL1A2, COL3A1. B. Top 50 genes from fibroblast signatures shows enrichment in PML and OSCC groups. C. Association of p-EMT and fibroblasts GSVA scores.

**S3.**
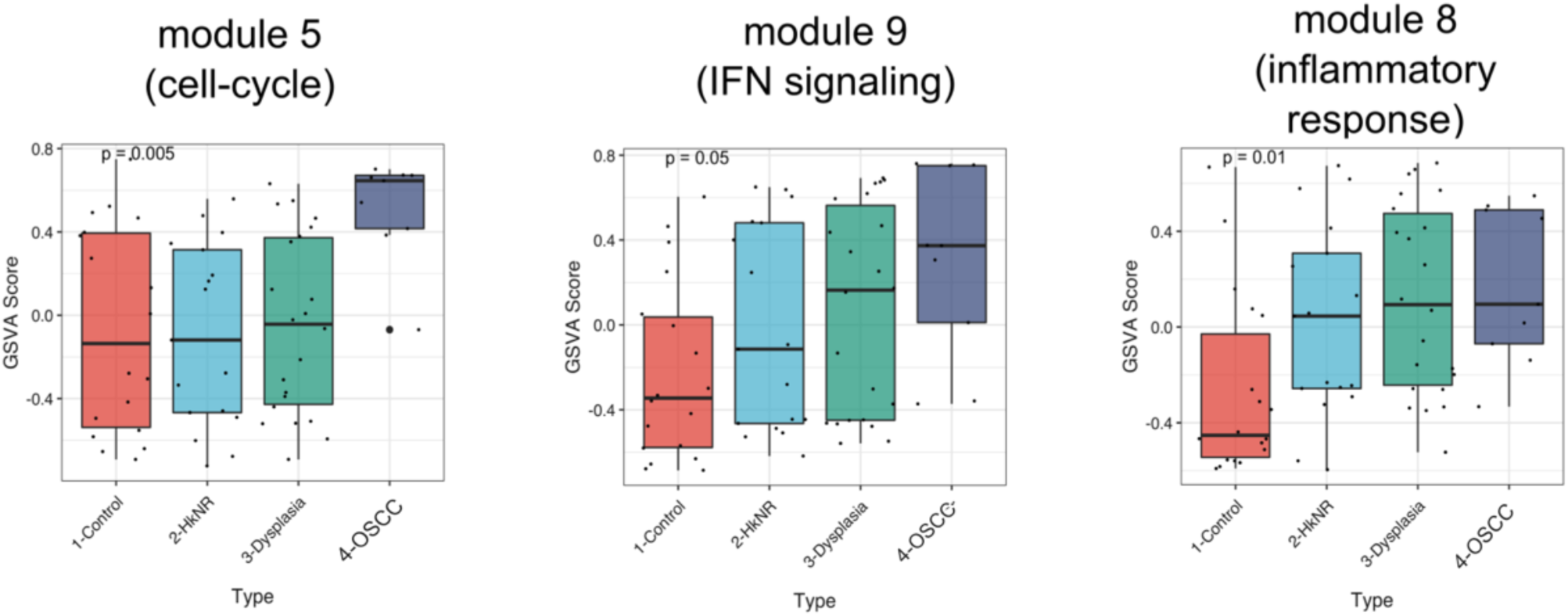
Enrichment of lung bronchus modules. GSVA enrichment scores from modules 5, 8, and 9 from lung bronchus PML study.

**S4.**
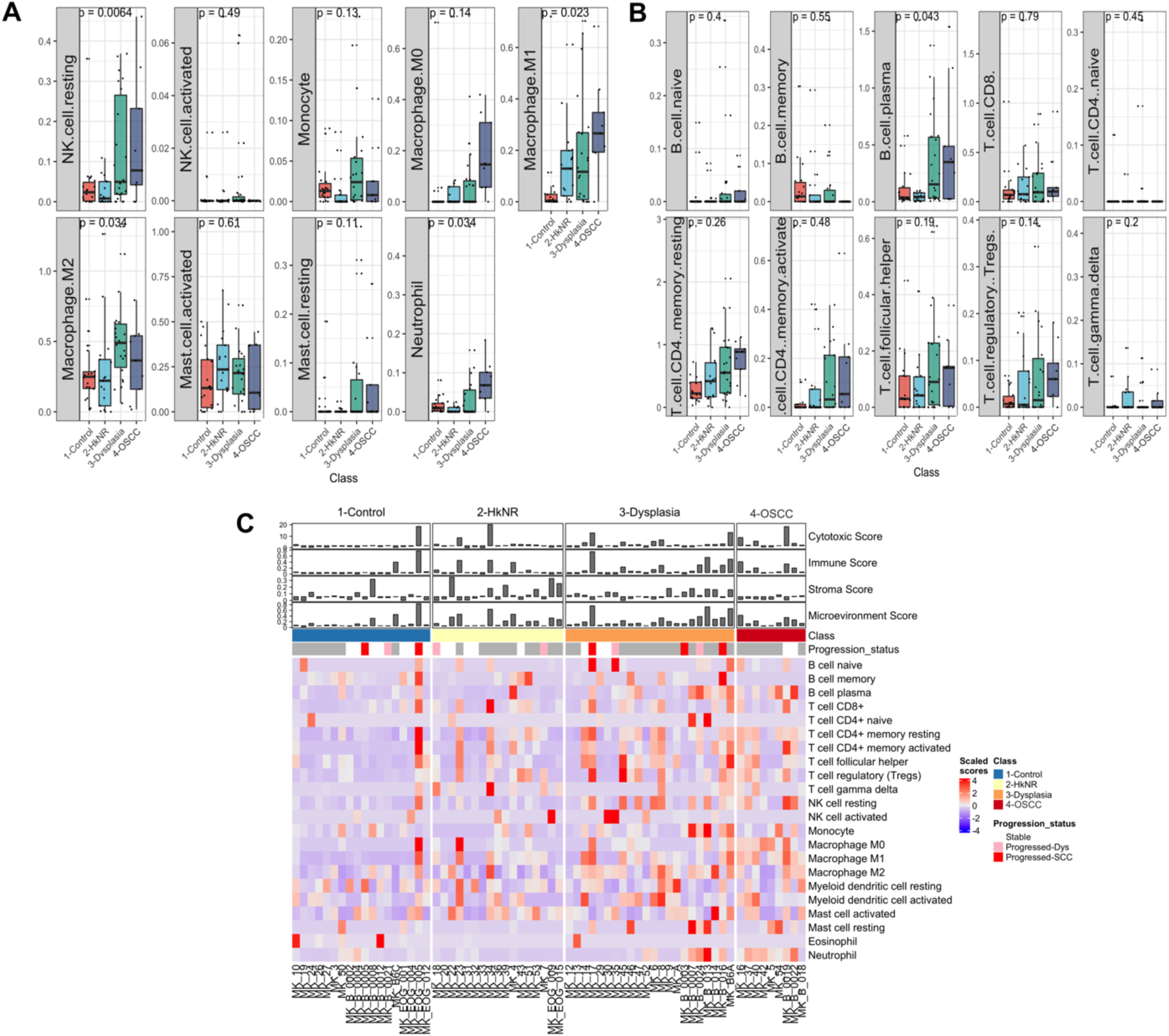
Cell-type deconvolution scores of immune sub-types A. innate, B. adaptive types. C. heatmap of abundances stratified by histopathology along with smoking and progression statuses.

**S5.**
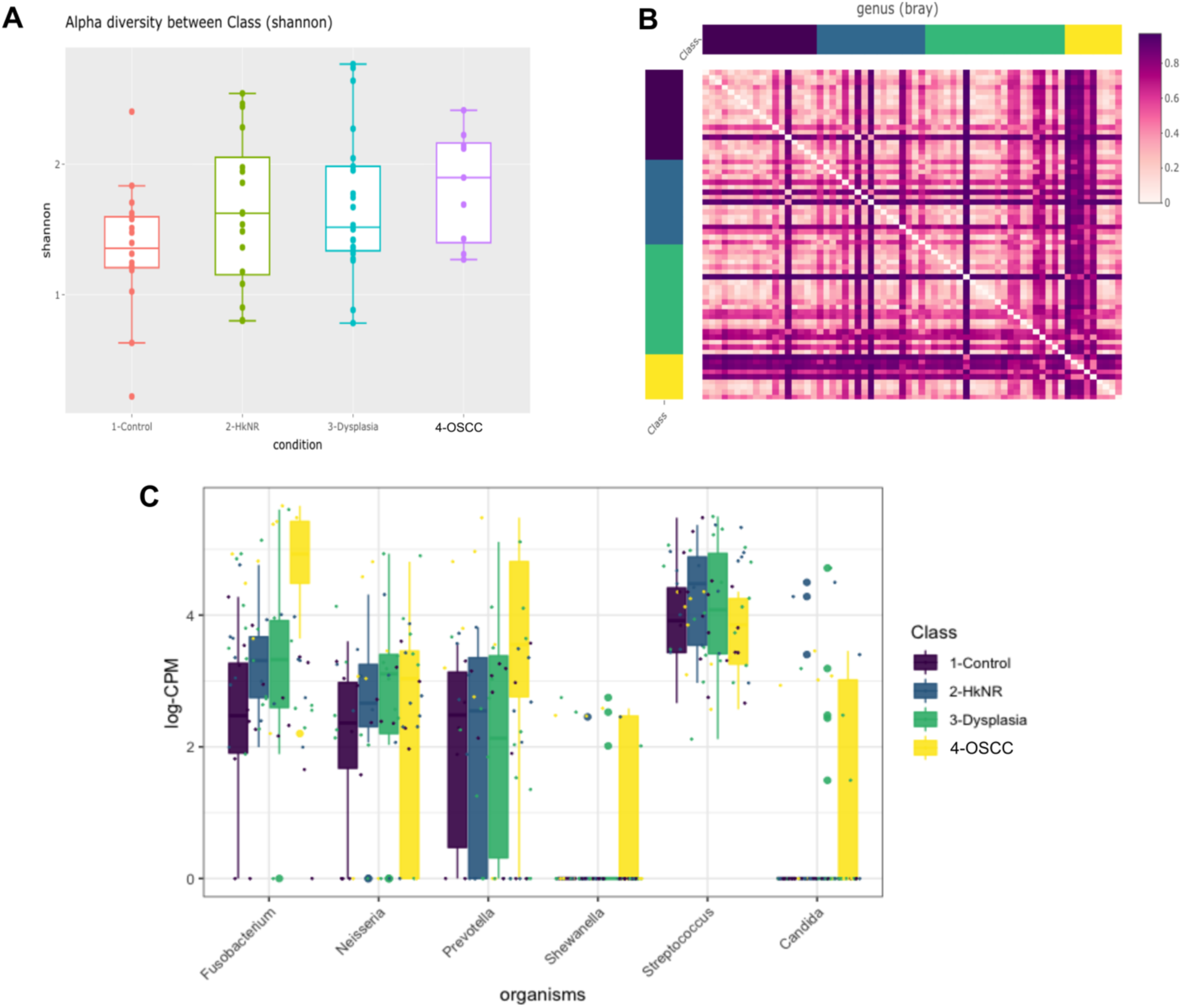
Microbial diversity analysis. A. Alpha diversity stratified by histopathological groups. B. Beta diversity stratified by histopathological groups. C. relative abundance in logCPM counts for selected disease-causing genera.

## Tables

Table 1. Patient summary by histopathology and clinical factors. Biopsy from patients (n=66) are separated into five histopathology groups and segregated by age, smoking and disease progression status.

## Supplementary Tables

ST1. Differential expression analysis results from DESeq2 for pairwise comparisons between the histopathological groups.

ST2. Pathway enrichment analysis of differentially expressed genes pairwise using hyper enrichment analysis using Hallmark compendium.

ST3. Pathway enrichment analysis of differentially expressed genes pairwise using hyper enrichment analysis using Reactome compendium.

ST4. Microbial differential abundant analysis results from DESeq2 for pairwise comparisons between the histopathological groups.

ST5. MSEA results of microbe-set associations with genes.

ST6. Hyper enrichment analysis of microbe-set genes from MSEA on enriched pathways from host analysis.

## Notes

### Competing Interest Statement

The authors have declared no competing interest.

